# Genetic associations of dimensional autistic phenotypes

**DOI:** 10.1101/2023.07.18.549458

**Authors:** Tore Eriksson, Chiaki Nakamori, Kazunari Iwamoto

## Abstract

Since there is a large variation in the symptoms shown by persons affected with ASD, analyzing genetics data using a case-control design is not straightforward. To avoid the difficult problem of defining heterogeneous groups, we used four different methods to compute a latent representation of a merged set of three psychometric tests. Computing the genetic contribution of each representation using a subset of participants with genetic data, we showed that factor analysis as well as variable autoencoders separates information contained in psychometric tests into genetically distinct phenotypic domains. Using the individual-level loadings of the domains as quantitative phenotypes in genome-wide association studies we detected statistically significant genetic associations in the domain related to insistence on routine, as well as suggestive genetic signals in other domains. We hope that these results can suggest possible domain-specific interventions in the future.

## Introduction

As its name suggests, one important aspect of autism spectrum disorder (ASD) is the heterogeneous nature of its symptoms. Some individuals display multiple symptoms and some only a subset. The severity of each symptom also varies among individuals. ASD is thus quantitative rather than qualitative, which implies that the case-control study is not an optimal strategy to analyze this disorder. Current diagnostic scales are thought to represent a combinations of constructs. Previous research has shown that CASI-4R scores could be mapped to a three-dimensional space—social interaction, communication, and repetitive behavior^1^.

We propose to combine dimensional analysis of phenotypes with genome-wide association studies. In this research we have attempted to extract latent dimensions from a combined set of psychometric test results. We have tried both the standard method latent variable modelling^2^, as well as dimensional reduction using deep learning models such as variational autoencoders with a low-dimensional latent space^3^. The clinical meaning of the found dimensions were elucidated by looking at the mappings of the original phenotypic data onto the latent space. As some of these dimensions are genetic while others will be related to social and environmental influences, we examined the genetic heritability of each concept.

Specifically, we selected a subset of subjects and psychometric tests with high coverage. The results were collected into matrix form and missing phenotype values were imputed. Latent variable modelling using this matrix produces a number of latent dimensions. Using the test results, individual subjects were mapped onto these dimensions. Genome-wide association studies (GWAS) using the position of the patients on these dimensions as independent quantitative traits were run using genetic variation data collected by the SPARK^4^ study. This research is expected to uncover genetic associations with behavioral traits that would be difficult to detect by an association study on the raw psychometric results directly. This could possibly lead to potential drug targets aimed not at ASD in general but at alleviating specific traits such as sensory hypersensitivity or repetitive body movements.

## Methods

### Participant selection

Initial data from SPARK (v8)^4^ contain data for 173,143 subjects after removing subjects that have withdrawn from the study. We further choose to exclude subjects with known large copy number variations (CNVs) or karyotype variations by inspection of the `gen_dx1_self_report` field. Excluded entries are listed in Supp. Table 1.

### Phenotype imputation

As phenotypes, we selected the Repetitive Behavior Scale-Revised (RBS-R), the Social-Communication Questionnaire-Lifetime (SCQ-L), and the Developmental Coordination Disorder Questionnaire (DCDQ). We removed subjects with a large number of unanswered questions and kept subject with results for all three tests. Remaining missing data were imputed using predictive mean matching as implemented in the mice package^5^.

### Latent factor models

Parallel analysis of factor scree values (R psych package^6^) implied the existence of six latent factors, but we decided to run our analysis considering eight factors to accommodate for information not present in the scree values. We tried four different methods to compute latent factors, two standard numerical methods and two based on deep learning.

### Numerical models

Principal component analysis (PCA) using scaled and centered values was used as the baseline method. As a standard psychometric method, factor analysis was run with maximum likelihood and oblimin rotation.

### Deep learning-based models

To allow for additional non-linearity we also built two different models based on deep learning techniques—a variational autoencoder and a transformer-inspired model. Both models had a latent space of size eight to make the results comparable to the latent factor model.

The variational autoencoder had two fully connected layers and a bottleneck layer to create en embedding of dimension eight (VAE1-8) for encoding, and two fully connected dense layers for decoding. The complete model definition is described in the supplementary information. Training was done on the imputed dataset with ordinal data mapped onto a variation of one-hot encoding where the response as well as lower values were mapped to one, excluding the lowest value. Training was done using the Adam optimizer with a VAE-specific loss function. A batch size of 32 was used with 500 epochs of training. 1000 samples were held back as an evaluation data set.

The transformer-inspired model was constructed with a separate embedding for each question. Since training with masking is robust even when missing values are present, this model was trained on the unimputed dataset. The model has a hidden space of dimension 256 and eight separate heads. The output of the start token was mapped to a vector of dimension eight using a fully connected layer and this vector was used to train a separate decoder with the same dimensions. Training was dome using the Adam optimizer and sparse categorical accuracy was used to compute accuracy of both encoder and decoder.

### Genome data QC and imputation

Genotypes were obtained from SPARK release iWES_v1.2022_02. We extracted data for the participants selected in our phenotyping analysis. For genomic quality control, PLINK (v1.9)^7^ was used. We first excluded subjects matching any of the following criteria: 1) ambiguous sex, 2) call rate below 90%, 3) heterozygosity outlier (mean ± 3 s.d.). Next, we excluded SNPs with call rate below 99%, minor allele frequency (MAF) < 0.01 or a Hardy–Weinberg proportion test with p > 10^−6^ from whole genome imputation. Imputation was performed based on 1000 Genomes Phase 3 data (GRCh38) as the reference panel using SHAPEIT4 (v4.1.3)^8^ and Beagle (v5.4)^9^ for phasing and imputation, respectively. After imputation, SNPs with an imputation score above 0.7 or an allele frequency above 0.01 were selected for use in association tests.

### Heritability and genomic correlations

Analysis of heritability and genetic correlation was done using BOLT-REML (v2.4)^10^ applying standard settings. Subject’s sex, age in months and genotyping batch were used as covariates.

### Genome-wide associations

Association test were done using a mixed model based method, BOLT-LMM (v2.4)^10^. Based on instructions, SNPs with imputation score above 0.9 and a pairwise LD r^2^ less than 0.2 (`--indep-pairwise 50 2 0.2` in PLINK) were selected for model fitting. The remaining SNPs were included as imputed SNPs in the association test. As above, subject’s sex, age in months and genotyping batch were used as covariates.

### Colocalization

Colocalization between GWAS signals and eQTLs was computed using data from the GTEx project (v8)^11^. To reduce noise, only gene-tissue pairs having strongly associated SNPs (p > 5×10^−8^) were used in the analysis. Colocalization statistics were computed using the `coloc.abf` method provided in the coloc package (v5.1.0.1)^12^.

### Gene set analysis

Gene set analysis was run using the GSEA method as implemented in the clusterProfiler package (v4.4.4)^13^. Normally, in methods like GSEA one uses gene expression fold changes or their p-values for ranking, but there is no corresponding value for a colocalization analysis. Instead we choose to use the product of the colocalization posterior probabilitie (`PP.H4.abf`) and the posterior probability of the lead SNP (`SNP.PP.H4`). GSEA was run against the human subset of the Hallmark set of pathways^14^.

## Results

### Phenotypic data

After excluding subjects that had withdrawn from the project, there were 173,143 available subjects initially. First, 798 subjects with known karyotypic abnormalities or large copy number variants were removed. Out of 41,260 subjects with RBS-R responses 23 were removed due to having more than 12 missing answers. Similarly, out of 79,404 subjects with SCQ-L responses 459 persons had more than six missing answers and were excluded. Finally, of the 30,940 subjects with DCDQ results, five with more than six missing answers were also removed. Combining these three sets, we ended up with 30,547 subjects who had reported data for all three scales (Fig. 1).

**Figure 1:**
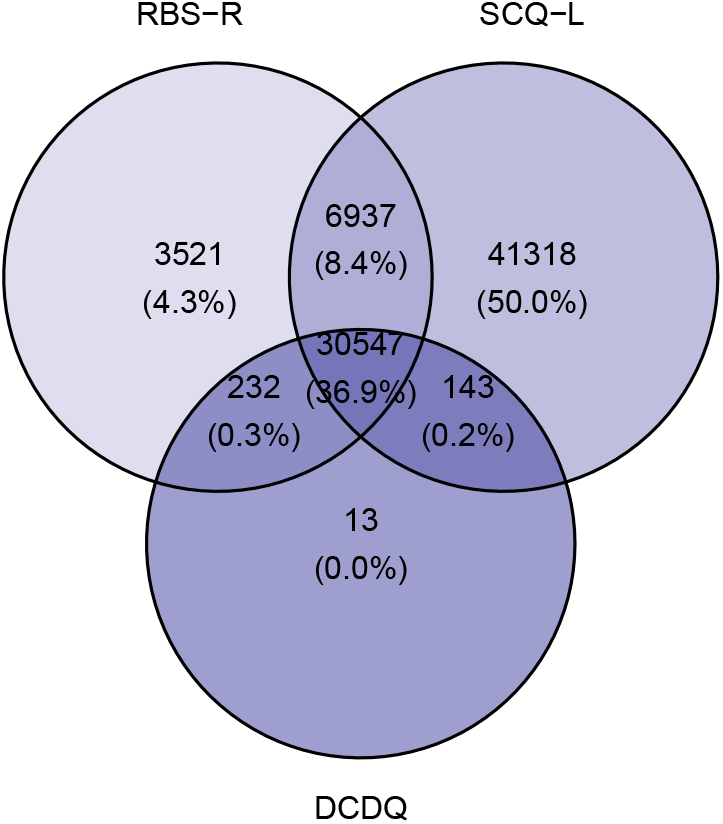
Venn diagram of subject overlaps.

Of our subjects, 6592 (22%) were female and 23,955 (78%) were male. The respective age distributions are shown in Fig. 2. The final phenotypic dataset contained 7,963 missing values which were imputed using the combined values from all three scales.

**Figure 2:**
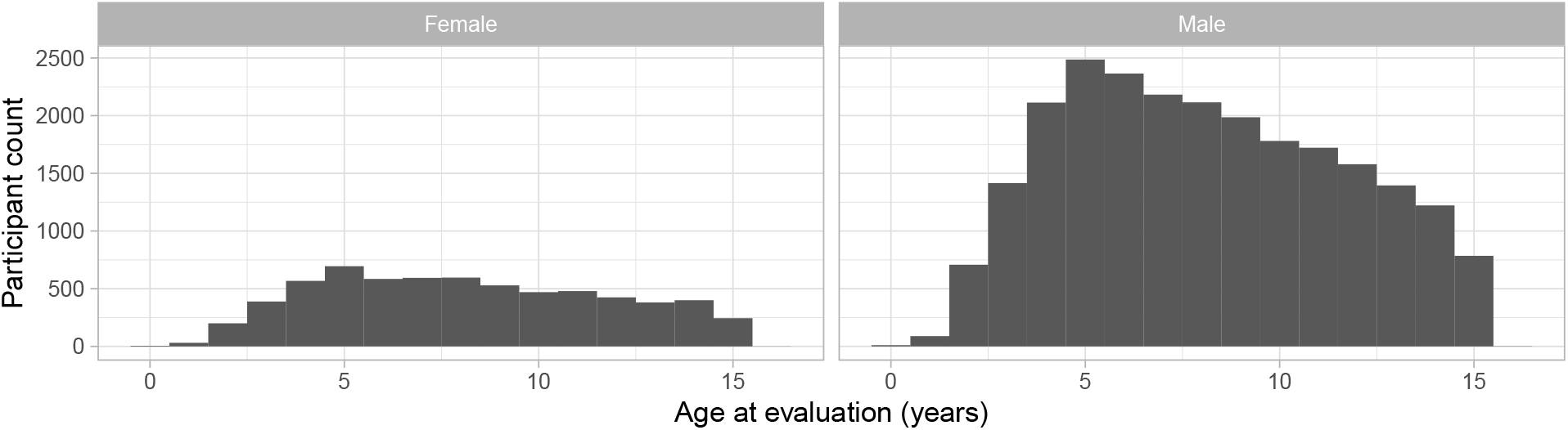
Distribution of participant age by sex.

### Latent factors

#### Principal Component Analysis

The top eight principal components (PC) explained 42% of the total variability (Fig. 3). All scores displayed a unimodal distribution (Fig. 4). There were no apparent sex-specific differences (Supp. Figure 1).

**Figure 3:**
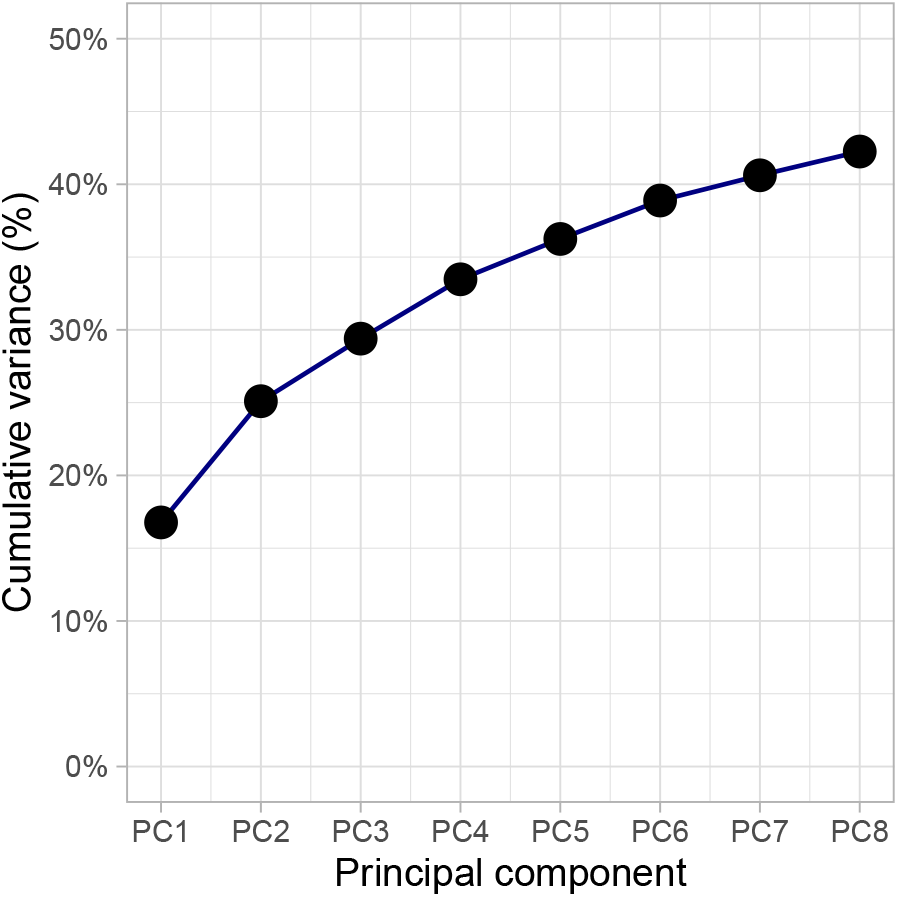
Cumulative percentage of PC explained variance.

**Figure 4:**
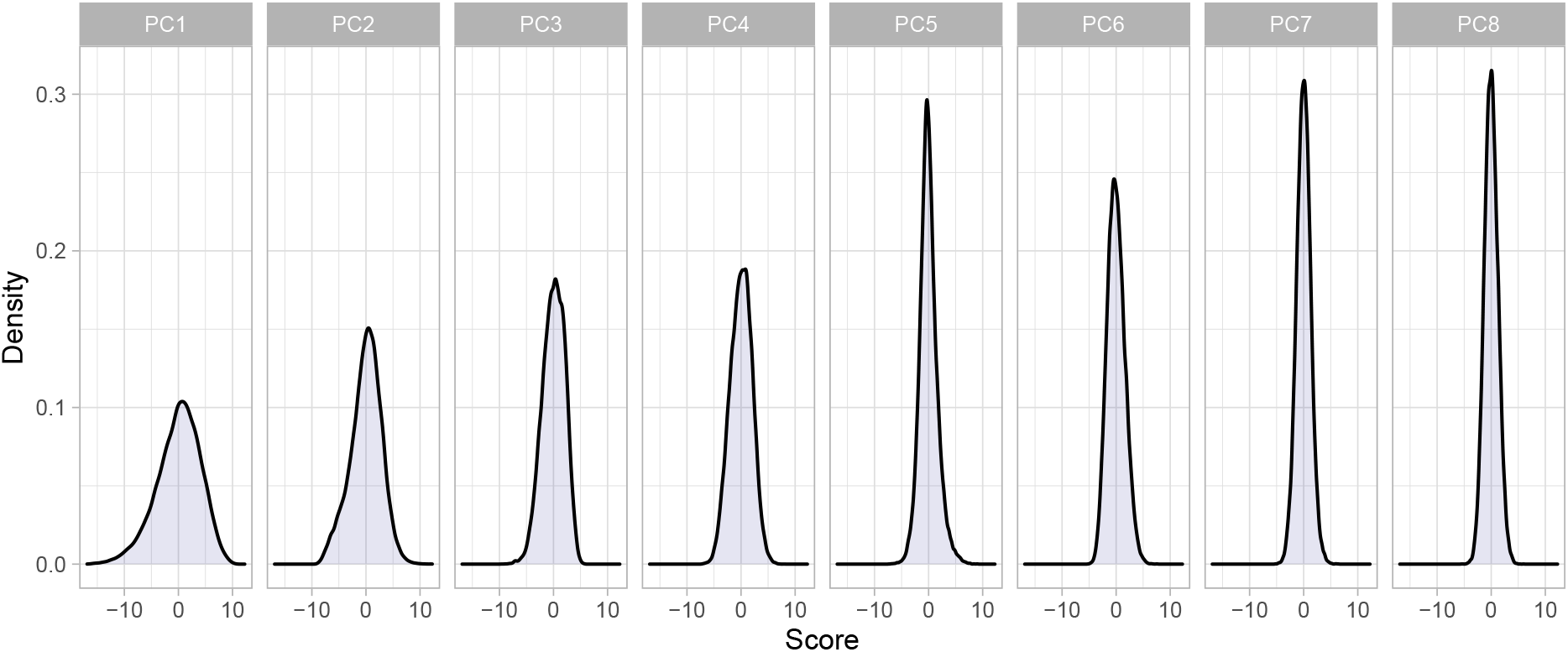
PC score distribution.

#### Factor analysis

The top eight latent factors (FA) explained 37% of the total variability (Fig. 5). Bimodal distributions were observed for FA2 and FA6, ant the other factors displayed unimodal distributions (Fig. 6). There were no apparent sex-specific differences (Supp. Figure 2).

**Figure 5:**
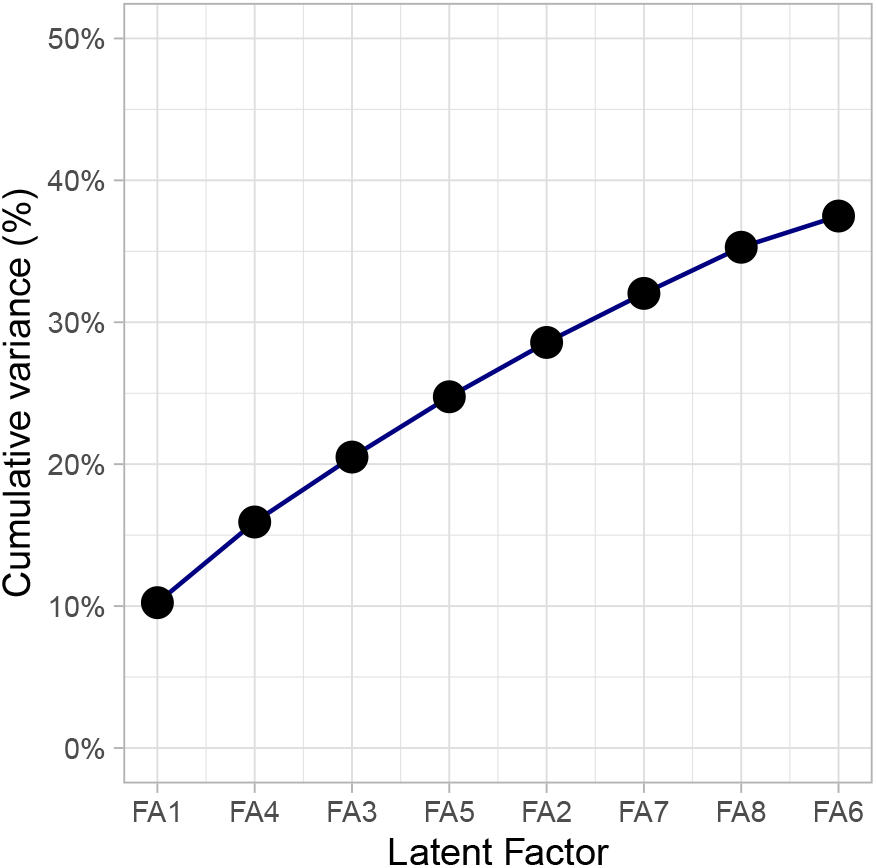
Cumulative percentage of FA explained variance.

**Figure 6:**
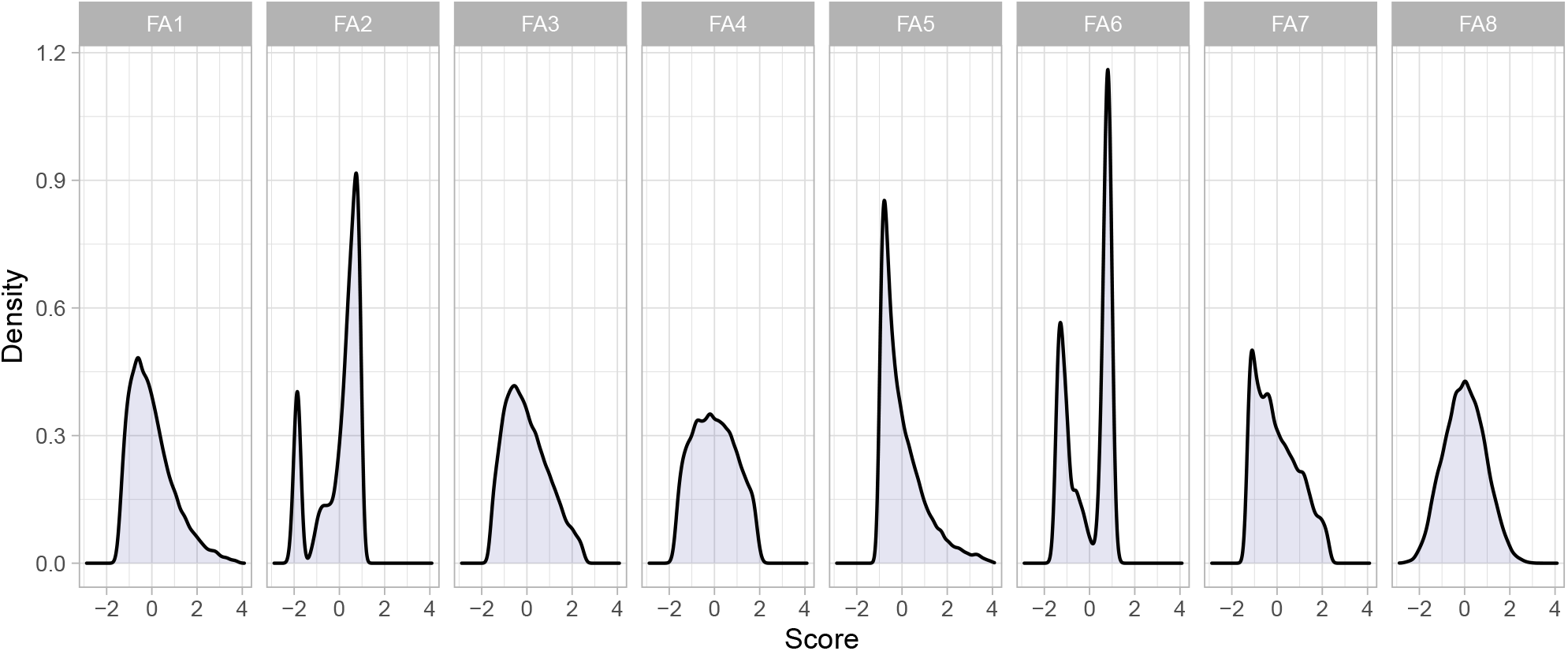
FA score distribution.

#### Deep learning-based models

The variational autoencoder reached a binary accuracy of 0.84 on the training data and 0.85 on the validation subset. The transformer reached a final sparse categorical accuracy of 0.76 on the full dataset. As expected, both models had similar distributions of latent dimensions, and as above there were no sex-specific differences (Supp. Figures 3 & 4).

### Heritability and genetic correlation

We computed the heritability for all four sets of latent factors (Fig. 7). A top heritability of around 0.4 was achieved by factor analysis (FA) FA1 and variational autoencoder (VAE) VAE8. This heritability was comparable to that observed for the SCQ-L total score (Supp. Figure 5).

**Figure 7:**
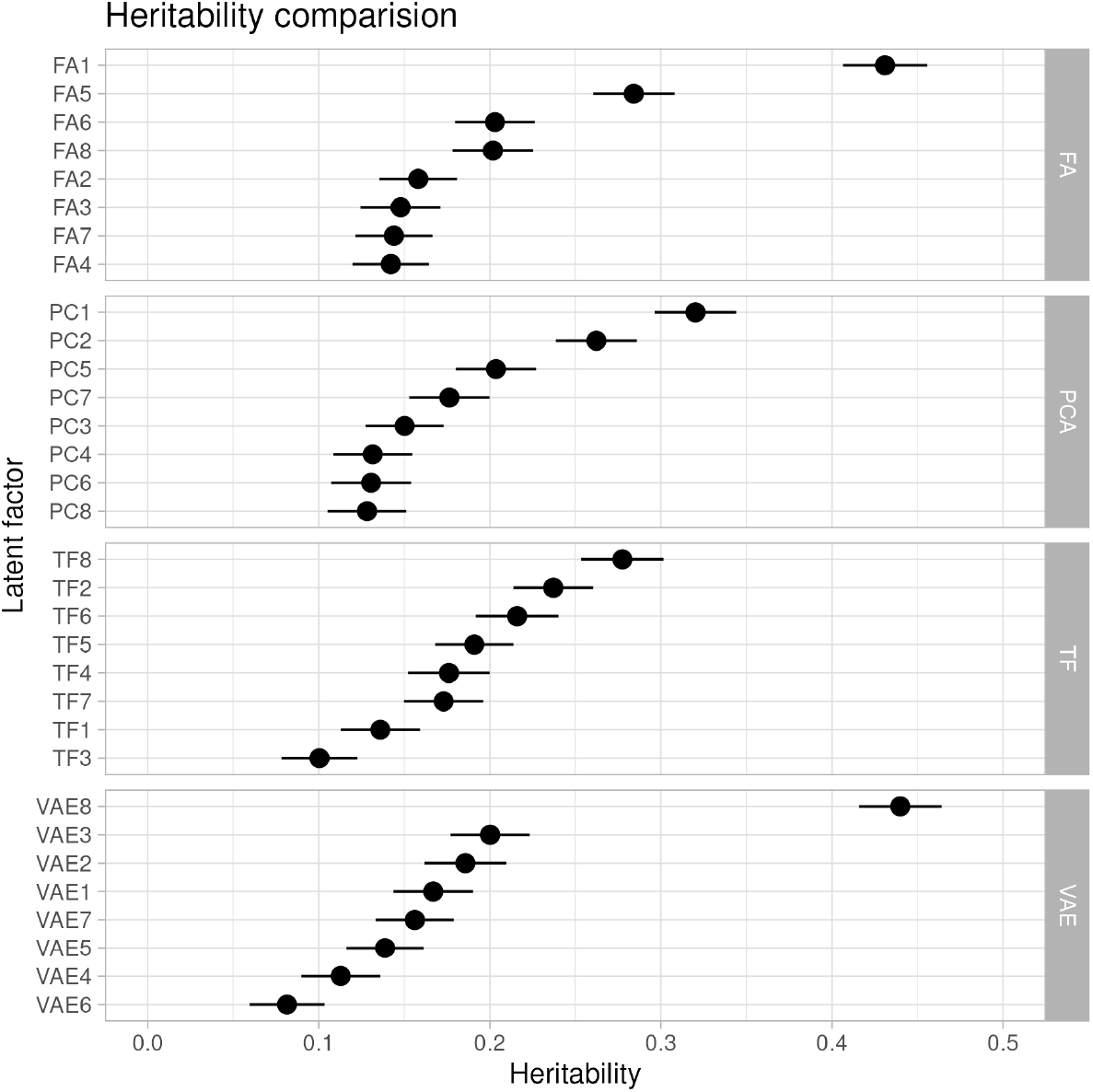
Latent factor heritability.

Looking at the result of clustering all factors by their genetic correlations (Supp. Figure 6), PCA factors displayed a tendency to have a lot of off-diagonal elements, which implies that this method is inconsistent with the other methods and unsuitable to separate the phenotypic signal. We also notice that the two deep learning models have learnt a number of factors not found by the numeric models. As the main factor for the transformer model showed a low heritability, we will henceforth focus on the results for the factor analysis and variational autoencoder models.

### Phenotypic domains

Studying the genetic correlation between the latent factors of our two main models (Fig. 8) as well as the numerical correlation between the latent factors and individual questions (Supp. Figures 7 & 8), we propose that the latent factors correspond to specific phenotypic domains shown in Table 1. The domain showing the strongest heritability (P1) is clearly involved with insistence on routine. Two domains related to speech (P2) and head movements (P6) are separate in factor analysis with a slight genetic correlation of 0.19, but the VAE merges them into one single latent factor (VAE3).

**Table 1:** Proposed domains. The main domain for VAE3 was P2, but it was also correlated to the P6 domain. Domains P8 to P10 were method-specific.

| Domain | Phenotype | FA | FA h | VAE | VAE h | FA-VAE corr. |
| --- | --- | --- | --- | --- | --- | --- |
| P1 | Routine | FA1 | 0.43 | VAE8 | 0.44 | 0.98 |
| P2 | Speech | FA2 | 0.16 | VAE3 | 0.2 | 0.69 |
| P3 | Gross motor skills | FA3 | 0.15 | VAE7 | 0.16 | -0.91 |
| P4 | Social interaction | FA4 | 0.14 | VAE5 | 0.14 | 0.81 |
| P5 | Self-harm | FA5 | 0.28 | VAE1 | 0.17 | -0.66 |
| P6 | Head movements | FA6 | 0.2 | VAE3 | 0.2 | 0.67 |
| P7 | Fine motor skills | FA7 | 0.14 | VAE6 | 0.08 | 0.45 |
| P8 | Communication | FA8 | 0.2 |  |  |  |
| P9 | Restricted interests |  |  | VAE2 | 0.19 |  |
| P10 | Bull in china shop |  |  | VAE4 | 0.11 |  |

**Figure 8:**
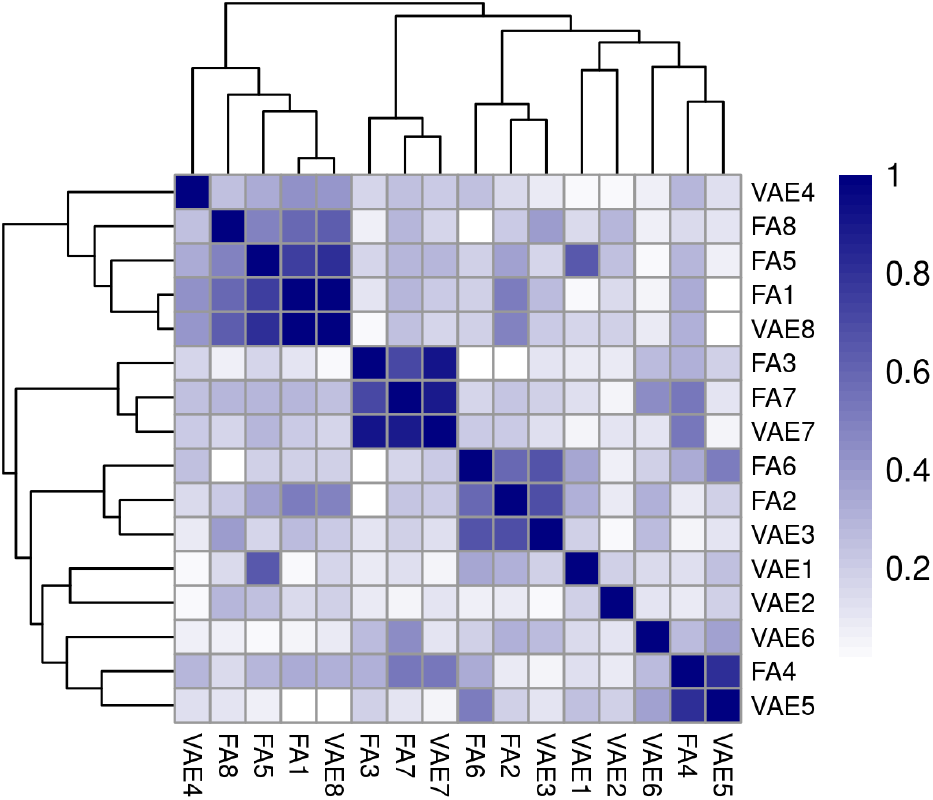
Latent factor genetic correlation between FA and VAE.

### Genome-wide associations

Genome-wide association results for the factor analysis results and the variational autoencoder latent dimensions for the 16,293 individuals with genetic data is shown in Fig. 9 and Fig. 10, respectively. 47 SNPs reached a p-value below the standard threshold of 5×10^−8^ (Sup. Table 2). A majority (35) of significant SNPs were from the latent factor FA1 which was the domain displaying the largest genetic heritability.

**Figure 9:**
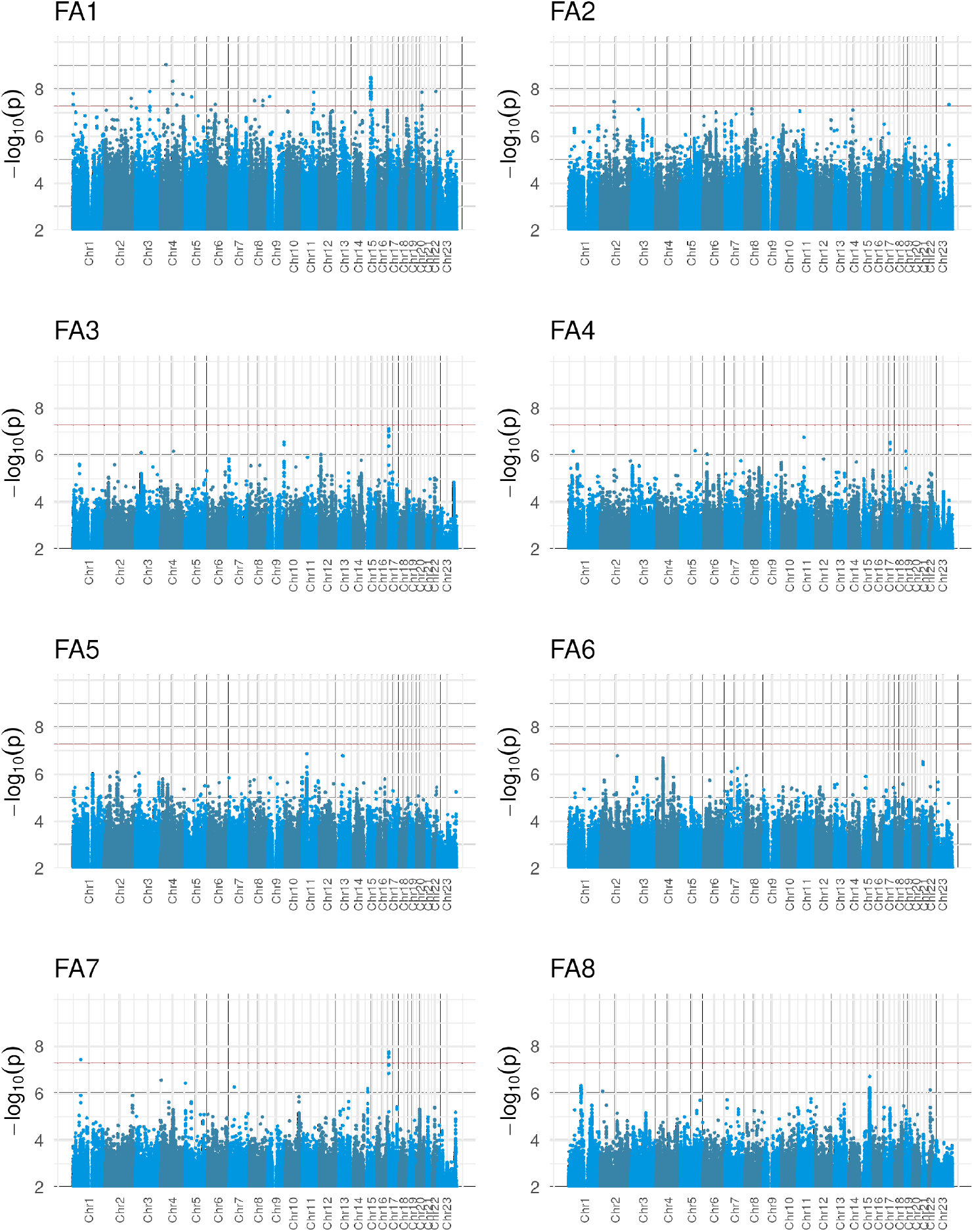
FA GWAS. Manhattan plot for GWAS based on latent factors. The vertical red line denotes a p-value of 5×10^−8^.

**Figure 10:**
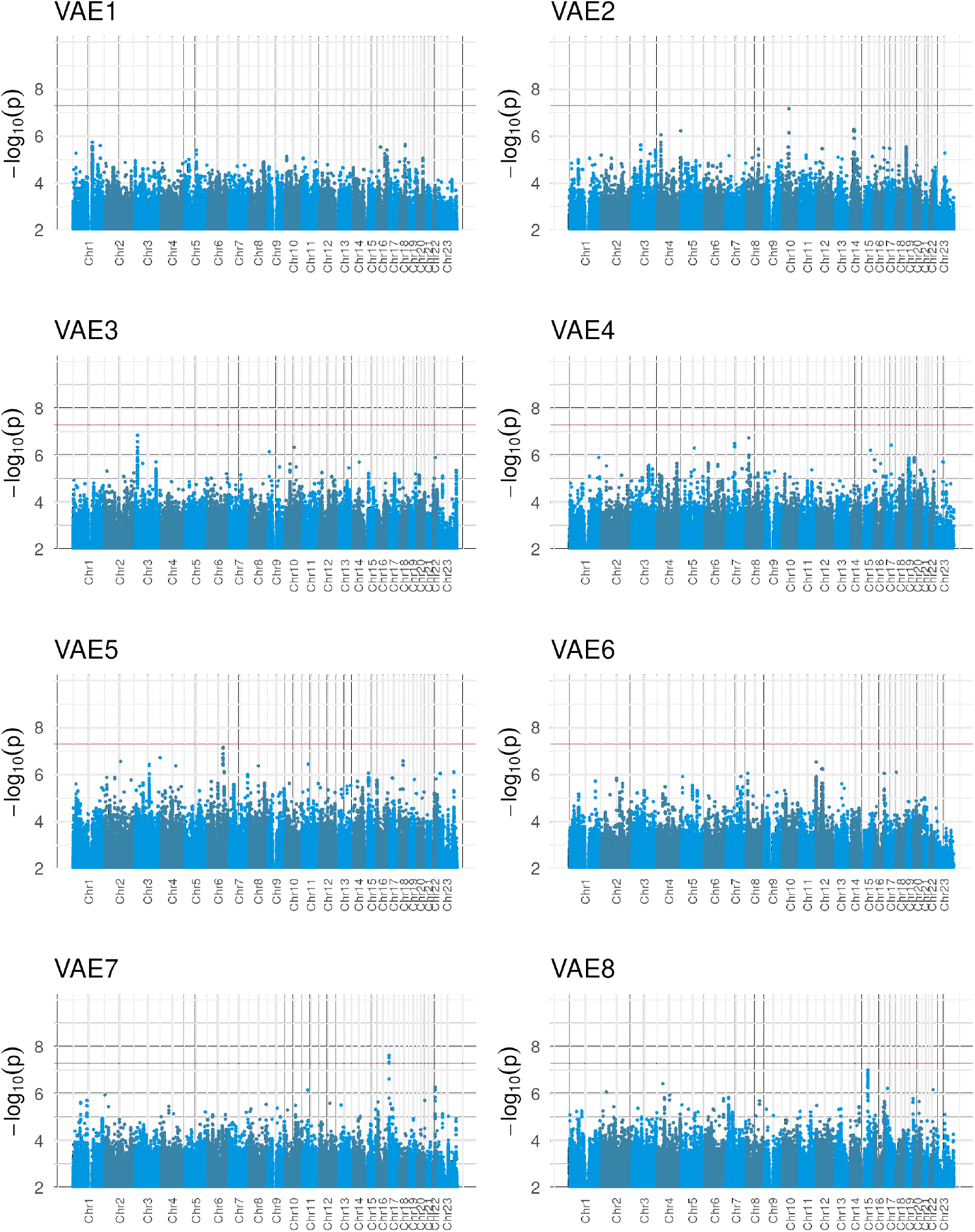
VAE GWAS. Manhattan plot for GWAS based on VAE latent dimension. The vertical red line denotes a p-value of 5×10^−8^.

Through manual inspection of genomic regions containing significant SNPs, we defined twenty different regions as having a clear genetic signal. By referring eQTL and coaccessibility data (not shown) as well as previous reports of involvement with ASD, we also selected one or two genes in each region as implicated genes (Table 2).

**Table 2:** Significant regions.

| Latent fact. | Chr. | Region<sup>*</sup> | Lead SNP | P-value | abs(Beta) | Implicated gene |
| --- | --- | --- | --- | --- | --- | --- |
| FA1 | 1 | 3.49 - 3.58 | 1_3553817_T_C | $1.5 \times 10^{-8}$ | $0.15 \pm 0.03$ | *MEGF6* |
| FA7 | 1 | 60.79 - 60.90 | 1_60797625_C_T | $3.6 \times 10^{-8}$ | $0.12 \pm 0.02$ | *NFIA* |
| FA2 | 2 | 109.43 - 109.48 | 2_109474702_G_A | $3.3 \times 10^{-8}$ | $0.24 \pm 0.04$ | *SH3RF3* |
| FA1 | 2 | 210.70 - 210.98 | 2_210974349_G_T | $2.4 \times 10^{-8}$ | $0.22 \pm 0.04$ | *ERBB4* |

| Latent fact. | Chr. | Region* | Lead SNP | P-value | abs(Beta) | Implicated gene |
| --- | --- | --- | --- | --- | --- | --- |
| FA1 | 3 | 114.73 - 115.23 | 3_114738193_T_C | $1.2 \times 10^{-8}$ | $0.33 \pm 0.06$ | *ZBTB20* |
| FA1 | 4 | 129.28 - 129.41 | 4_129413093_TTG_T | $4.7 \times 10^{-8}$ | $0.07 \pm 0.01$ | *PCDH10* |
| FA1 | 4 | 177.40 - 177.80 | 4_177571470_A_G | $1.6 \times 10^{-8}$ | $0.25 \pm 0.04$ | *NEIL3* |
| FA1 | 4 | 45.57 - 46.17 | 4_46167100_GC_G | $8.6 \times 10^{-10}$ | $0.12 \pm 0.02$ | *GABRG1/GABRA2* |
| FA1 | 4 | 99.20 - 99.32 | 4_99322288_T_G | $4.4 \times 10^{-9}$ | $0.07 \pm 0.01$ | *ADH1B* |
| FA1 | 5 | 57.69 - 57.83 | 5_57755770_C_G | $2.1 \times 10^{-8}$ | $0.07 \pm 0.01$ | *PLK2* |
| FA1 | 6 | 62.84 - 63.18 | 6_62858830_CA_C | $4.3 \times 10^{-8}$ | $0.07 \pm 0.01$ | *PHF3* |
| FA1 | 8 | 106.94 - 107.36 | 8_107065417_T_C | $3.0 \times 10^{-8}$ | $0.35 \pm 0.06$ | *ANGPT1* |
| FA1 | 8 | 41.63 - 41.86 | 8_41855161_G_A | $3.0 \times 10^{-8}$ | $0.23 \pm 0.04$ | *ANK1* |
| FA1 | 9 | 15.11 - 15.16 | 9_15148270_C_CT | $2.0 \times 10^{-8}$ | $0.15 \pm 0.03$ | *ZDHHC21* |
| FA1 | 11 | 93.28 - 93.44 | 11_93326188_T_C | $1.3 \times 10^{-8}$ | $0.29 \pm 0.05$ | *DEUP1* |
| FA1 | 15 | 53.68 - 53.76 | 15_53691652_T_C | $3.1 \times 10^{-9}$ | $0.08 \pm 0.01$ | *UNC13C* |
| FA7 | 17 | 3.20 - 3.49 | 17_3275656_G_A | $1.7 \times 10^{-8}$ | $0.17 \pm 0.03$ | *OR1E1* |
| FA1 | 20 | 42.73 - 42.74 | 20_42735872_C_T | $1.3 \times 10^{-8}$ | $0.08 \pm 0.01$ | *PTPRT* |
| FA1 | 22 | 41.20 (single) | 22_41202352_T_C | $1.2 \times 10^{-8}$ | $0.56 \pm 0.10$ | *EP300* |
| FA2 | X | 129.80 - 129.86 | 23_129859450_T_C | $4.4 \times 10^{-8}$ | $0.20 \pm 0.04$ | *ZDHHC9* |

\* hg38 coordinates (Mbp)

The strongest peak (FA1 P = 8.6×10^−10^) is located on chromosome four, in the intergenic region between *GABRG1* and *GABRA2*. Both genes code for components of the GABA_A_ receptor. This receptor has been implicated in the etiology of ASD for many years^15,16^.

The second strongest peak (FA1 P = 3.1×10^−9^) is within the *WDR72* gene. While we could not find any compelling link to autism for this gene, this peak is also located 140 kbp upstream of a transcription start site of the gene *UNC13C*. This gene is expressed in brain, predominantly in inhibitory neurons^17^, and copy number variants including this gene has also been implicated in attention-deficit hyperactivity disorder^18^. Since the family of UNC13 proteins are known to regulate neurotransmitter release at synapses^19^, we believe that *UNC13C* is a strong candidate for this peak.

The third strongest peak (FA1 P = 4.4×10^−9^) was found in the alcohol dehydrogenase cluster on chromosome four, implicating the promoter region of the gene coding for alcohol dehydrogenase 1B, *ADH1B*. This cluster of alcohol dehydrogenase genes have been linked to autism in previous studies as well^20^. Although the SNP is located close to the transcription start site of *ADH1B*, it is also reported to be an adipose tissue eQTL for the *ADH1A* gene.

All these three peaks were associated with the P1 phenotype, which was the phenotype displaying the largest genetic heritability. A majority of the other significant peaks were found in phenotype P1 as well (Table 2). Genes with a high confidence on involvement in ASD according to the SFARI Gene^21^ database included *PHF3, EP300* (both SFARI tier 1), *PCDH10*, and *PTPRT* (both tier 2).

### Colocalization

The colocalization study using eQTL data from the GTEx project^11^ detected an average of 191 genes for the factor analysis model and 129 genes for the variational autoencoder. We also examined which tissues had the largest ratio of colocalized genes for each latent factor (Table 3). There was no clear enrichment for brain regions among the top tissues, and we also observed a lack of concordance between the corresponding factors of the respective models.

**Table 3:** Tissues having colocalized genes. The percentages show the ratio of colocalized genes with a posterior probability (`PP.H4.abf`) > 0.5 to the number of all genes with significant eQTLs in each tissue.

| Phenotype | Factor | First tissue | Second tissue |
| --- | --- | --- | --- |
| P1 | FA1 | EBV-transformed lymphocytes (0.88%) | Stomach (0.87%) |
| P1 | VAE8 | EBV-transformed lymphocytes (0.29%) | Esophagus Mucosa (0.27%) |
| P2 | FA2 | Small Intestine Terminal Ileum (0.76%) | Brain Cerebellar Hemisphere (0.74%) |
| P2 | VAE3 | Kidney Cortex (0.38%) | Minor Salivary Gland (0.37%) |
| P3 | FA3 | Brain Hypothalamus (0.4%) | Spleen (0.34%) |
| P3 | VAE7 | Uterus (0.29%) | Vagina (0.23%) |
| P4 | FA4 | Spleen (0.36%) | EBV-transformed lymphocytes (0.34%) |
| P4 | VAE5 | Colon Transverse (0.47%) | Stomach (0.41%) |
| P5 | FA5 | Kidney Cortex (0.38%) | Cultured fibroblasts (0.31%) |
| P5 | VAE1 | Liver (0.31%) | Heart Left Ventricle (0.29%) |
| P6 | FA6 | Whole Blood (0.46%) | Spleen (0.44%) |
| P7 | FA7 | Liver (0.31%) | Vagina (0.3%) |
| P7 | VAE6 | Uterus (0.36%) | Artery Tibial (0.31%) |
| P8 | FA8 | Kidney Cortex (0.57%) | Liver (0.35%) |
| P9 | VAE2 | Brain Amygdala (0.35%) | Brain Hippocampus (0.27%) |
| P10 | VAE4 | Brain Substantia nigra (0.55%) | Brain Hippocampus (0.5%) |

### Hypothesis-free gene set analysis

Gene set enrichment analysis of genes ranked by the posterior probability of the top SNP in the colocalization analysis did not detect any significant gene categories when correcting for multiple hypothesis testing. Categories with an uncorrected p-value below 0.05 are listed in Supp. Tables 1 to 8.

## Discussion

Genetic causes of ASD are believed to be mostly rare single nucleotide and copy number variants which are difficult to detect using case-control studies. However, the hypothesis of this research is that the difference in *symptoms* displayed by ASD subjects could be modulated by common variants, and would thus be amendable for genome-wide association studies.

Previous research using a similar approach tried to separate ASD subjects into phenotypical subgroups and perform a case-control GWAS for each subgroup^22^. However, this approach does not match the current view of ASD as, as inherent in its name, a *spectrum disorder*. We thus propose that it is necessary to look at ASD phenotypes as quantitative entities.

Another approach is to use scores from psychometric test as quantitative measures^23^. Although psychometric tests used for diagnosis are by their nature numeric, they are not optimal for phenotyping since they use ordinal scales which are not inherently linear. Also, different tests focus on specific aspects of how the disorder presents itself, and thus the coverage of phenotypic variability varies between test. To account for these factors, we used rich phenotypic data collected by the SPARK project^4^ and choose to work on a subset of participants that had complete results from three different test. To extract phenotypic dimensions contained in the test results, we evaluated four different methods for detecting latent dimensions from the three merged tests results, fixing the number of dimensions to eight for all methods.

Visual inspections of loadings (for PCA and FA) or test score correlations (for TF and VAE) showed that all methods were able to separate the phenotypic results into latent vectors that were broadly aligned with known aspects of clinical phenotypes like repetitive behavior, tendency to self-harm, and communication difficulties.

By using the genetics data available for 16,293 individuals, we evaluated the heritability of each factor and concluded that factor analysis and VAE were the most promising methods. The main difference between the methods is that factor analysis allows for correlation between separate factors, while the VAE, by design, minimizes the numerical correlation between the latent dimensions. Both methods had a maximal heritability of about 0.4, which is in the same order of that seen with the RBS-R scale (Supp. Fig. 5).

We expected that the large heritability shown would lead to clear hits in the following association study, but no more than 47 SNPs reached the genome-wide significance threshold of p < 5×10^−8^. However, manual inspection of gene regions with SNPs displaying low p-values show peaks that are specific for each latent factor. The plausibility of these peaks will have to be evaluated by looking at epigenetic data and eQTL colocalization analyses.

In order to elucidate the functional implication of each phenotype, we performed gene set analyses against the gene sets using the Hallmark set of pathways^14^. Colocalization analysis with GTEx eQTLs were used to map genetic signals to genes. We were unable to find any pathways that were significant after multiple hypothesis correction. Inspecting the top hits, one intriguing result was the implication of KRAS signalling in domain P1 (Supp. Tables 3 and 16), as the set of genetic diseases known as RASopathies have been reported to display traits related to ASD^24^. One possible reason for the lack of robust results of this analysis is that visual inspection of the respective peaks showed that many peaks were located in putative enhancer regions, which are insufficiently covered by the GTEx analysis since they are at a distance of more than 1 Mbp from transcription start site.

### GABA signalling

Due to the lack of significant results from our global analysis, we choose to evaluate the found associations manually as well. The top peaks were all found in the P1 (FA1) domain. The top two peaks implicated genes directly involved in inhibitory GABA signalling through the GABA_A_ receptor. Although the consequences of mutations in the receptor vary between reports, the receptor is thought to be involved in modulating excitatory and inhibitory (E/I) balance in the brain^16^.

The relevance of ethanol to autism suggested by the third strongest peak in the alcohol dehydrogenase gene cluster is also thought to be related to GABA signalling and the GABA_A_ receptor. Both the *GABRG1* locus and the *ADH1A*/*ADH1B* locus have also been linked to alcohol consumption and alcohol usage disorders^20,25,26^. Though ethanol has been shown to regulate the activity of GABA receptors directly^27^, the patient population used in this study is predominantly composed of children who we assume do not consume alcohol. The genetic relationship between ASD and alcohol suggested by these results could be an indirect result of alcohol consumption during pregnancy, but epidemiological studies of this topic did not show a clear relationship^28,29^. Further research could explore this in more detail by looking at whether the *ADHB1* alleles linked to ASD were inherited from the mother or not.

Another possible source of ethanol is gut microbiota. A recent study^30^ described increased levels of ethanol in the portal vein of individuals with non-alcoholic fatty liver disease (NAFLD). Furthermore, a metagenomic analysis also showed a correlation between microbiome composition and postprandial ethanol concentrations. The strongest positive correlation was with the *Lactobacillus* species, a species that is also reported to have increased abundance in subjects with ASD^31^. Combined with reports of an increased risk of NAFLD in people on the spectrum^32^, evaluating the link between gut ethanol production and GABA signalling could be a promising direction for further studies.

### Fragile X syndrome

At least 5% of ASD cases are caused by single-nucleotide variants. To avoid genetic variants displaying an overwhelming effect on the phenotype we elected to remove individuals with a diagnosis of monogenic disorders. However, we were interested in whether variants with a milder effect on gene function would be related to our phenotypes.

The gene causing Fragile X syndrome, *FMR1*, show a prominent peak in the P7 domain (FA7 p = 6.5×10^−6^) overlapping with eQTLs for the gene. The main phenotype that this component represents is fine motor skills, which is consistent with what is reported in Fragile X patients^33^. The top peak (FA7 p = 1.7×10^−8^) in this component is located in a cluster of olfactory receptors on chromosome 17. Surveying neighboring genes, this peak is 50 kbp downstream of the gene coding for the cation channel TRPV3. As this channel has been shown to be expressed in Purkinje fibers in the rat cerebellum, and as intra-cerebellar inhibition of TRPV3 signalling reduced motor coordination in rats^34^, we think TRPV3 is linked to this trait in humans as well.

### Self-harm

We were also interested in the domain P5 which was related to self-harm. That there is a need to *treat* traits related to *i*.*e*. social interaction is a question that is debated among people on the spectrum as they see this trait as a part of their neurodiversity^35,36^. However, we think that this is of less concern in the case of self-harm, since this is evidently harmful for the individual and that having a treatment would be beneficial. We were unable to find statistically significant genetic signals on this component in our study, but we could observe a number of suggestive (p < 5×10^−5^) peaks that we review below.

The second largest peak (FA5 p = 1.6×10^−7^) was located on chromosome 13, upstream of the gene *OLFM4*, a shared loci for major depressive disorder, insomnia and chronic pain^37^. Another strong signal (FA5 p = 9.6×10^−7^) was located on chromosome one close to *KCNN3*, a gene linked to migraine in multiple studies^38,39^. As migraine and depression are believed to share underlying etiology^40^, and as migraine has been linked to sensory hyperreactivity in children with ASD^41^, our results suggests that that self-harm could be related to the same mechanism. As it is difficult to ask toddlers and non-communicative individuals whether they experience migraines, possibly with auras, one approach might be to use non-invasive methods like electroencephalography (EEG) to check for evidence of migraine attacks during bouts of self harm.

## Conclusion

Our goal with this research was to map clinical data for people on the autism spectrum into a phenotypic space of an arbitrary dimensional size of eight. Having evaluated different methods to create this space, is seem like the standard method of factor analysis gives dimensions which contain the strongest genetic signal, but it is still unclear if the strength of this signal is due to the inability of this method to separate the space into uncorrelated components. From a biological perspective, it is not clear whether the components *should* be correlated or not and we hope these results will lead to further studies of this point.

Our methods were able to distinguish a phenotypic domain centered on insistence on routine that displayed strong heritability. Although no clear biological mechanism could be determined trough a hypothesis-free analysis of genes linked to this domain, manual inspection of the top peaks implied GABA signalling through the GABA_A_ receptor as an important factor.

The remaining phenotypic domains did not show a strong heritability, which could imply that they have predominantly environmental origins. However, looking at strongly associated regions we were able to discern a possible connection of self-harming behavior to genes known to be involved in pain and migraine. There have been suggestions that children displaying self-harming behavior like head banging have a decreased sensitivity for pain, but one could also imaging that this behavior is due to experiencing a persistent low-grade migraine.

We are aware that the lack of clear results from hypothesis-free methods in this study leaves us with an incomplete evaluation of a majority of our genome-wide results. However, we do believe that a dimensional model of phenotypes is a promising approach to genetic studies in diseases that show large variation in the strength of symptoms. An increasing amount of available phenotype data, especially consisting of quantitative measurements in combination with improvement in numerical methods, will surely allow for even better models in the near future.

## Limitations

Even though we didn’t do any explicit filtering on diagnosis, after selecting participants with complete data sets all remaining individuals were diagnosed with ASD. Since ASD is a spectrum disorder, we would have preferred to include a large amount of normotypic individuals into the analysis to more fully cover the total range of symptoms. Although we also removed subjects with a genetic diagnosis of monogenic disorders like Fragile X syndrome, some undiagnosed individuals might be included and this could reduce the power of the genetic association studies.

Another concern is that the neurophysiological tests we used are optimized for diagnosis, but not for phenotyping. The latent factor analysis used will consider the correlation between specific questions, but the inherit fuzziness of ordinal data makes them sub-optional to use as quantitative measurements. We would have preferred to work with phenotypic data measured directly, as for example data on eye tracking and facial avoidance and we hope that such data will be available in the future.

We are also concerned with the bias inherit in the tests deployed, since the test designers are already selecting questions that query a specific domain, which will make it impossible to extract information for other domains. It is our hope that combining multiple tests will have ameliorated this problem in this study.

## Author contributions

Conceptualization, TE; data curation, TE; investigation, TI and KE; methodology, TE, CN and KI; software, TE, CN and KI; visualization, TE and KI; writing, TE and KI; resources, TE; supervision., TE.

## Acknowlegments

We appreciate obtaining access to phenotypic and genetic data on SFARI Base^42^.

The Genotype-Tissue Expression (GTEx) Project was supported by the Common Fund of the Office of the Director of the National Institutes of Health, and by NCI, NHGRI, NHLBI, NIDA, NIMH, and NINDS. The data used for the analyses described in this manuscript were obtained from the GTEx Portal on May 11, 2021.

