## Supplementary material for "Genetic associations of dimensional autistic phenotypes": Supplemntary information

Supplementary information

June 30, 2023

### Supplementory information

#### Variational Autoencoder definition

#### Full VAE

**Bottleneck** `z_mean`
**Dense layer** `z_log_var`
**Lambda function** `z_mean + k_exp(z_log_var/2) * epsilon`

```
Model: "autoencoder_VAE"
____________________________________________________________________________________________________
Layer (type)                     Output Shape          Param #     Connected to
====================================================================================================
input_2 (InputLayer)             [(None, 229)]         0

____________________________________________________________________________________________________
E1 (Dense)                       (None, 128)           29440       input_2[0][0]
____________________________________________________________________________________________________
dropout_9 (Dropout)              (None, 128)           0           E1[0][0]
____________________________________________________________________________________________________
E2 (Dense)                       (None, 32)            4128        dropout_9[0][0]
____________________________________________________________________________________________________
dropout_8 (Dropout)              (None, 32)            0           E2[0][0]
____________________________________________________________________________________________________
bottleneck (Dense)               (None, 8)             264         dropout_8[0][0]
____________________________________________________________________________________________________
dense (Dense)                    (None, 8)             264         dropout_8[0][0]
____________________________________________________________________________________________________
concatenate (Concatenate)        (None, 16)            0           bottleneck[0][0]
                                                                   dense[0][0]
____________________________________________________________________________________________________
lambda (Lambda)                  (None, 8)             0           concatenate[0][0]
____________________________________________________________________________________________________
dense_1 (Dense)                  (None, 32)            288         lambda[0][0]
____________________________________________________________________________________________________
dropout_10 (Dropout)             (None, 32)            0           dense_1[0][0]
____________________________________________________________________________________________________
dense_2 (Dense)                  (None, 128)           4224        dropout_10[0][0]
____________________________________________________________________________________________________
dropout_11 (Dropout)             (None, 128)           0           dense_2[0][0]
____________________________________________________________________________________________________
output (Dense)                   (None, 229)           29541       dropout_11[0][0]
====================================================================================================
Total params: 68,149
Trainable params: 68,149
Non-trainable params: 0
____________________________________________________________________________________________________
```

#### Encoder part

```
Model: "encoder_VAE"
_________________________________________________________________________________________________
Layer (type)                             Output Shape                             Param #
=================================================================================================
input_2 (InputLayer)                     [(None, 229)]                            0
_________________________________________________________________________________________________
E1 (Dense)                               (None, 128)                              29440
_________________________________________________________________________________________________
dropout_9 (Dropout)                      (None, 128)                              0
_________________________________________________________________________________________________
E2 (Dense)                               (None, 32)                               4128
_________________________________________________________________________________________________
dropout_8 (Dropout)                      (None, 32)                               0
_________________________________________________________________________________________________
bottleneck (Dense)                       (None, 8)                                264
=================================================================================================
Total params: 33,832
Trainable params: 33,832
Non-trainable params: 0
_________________________________________________________________________________________________
```

### Supplementary Figures

#### Supp. Figure 1: PCA loadings by sex.

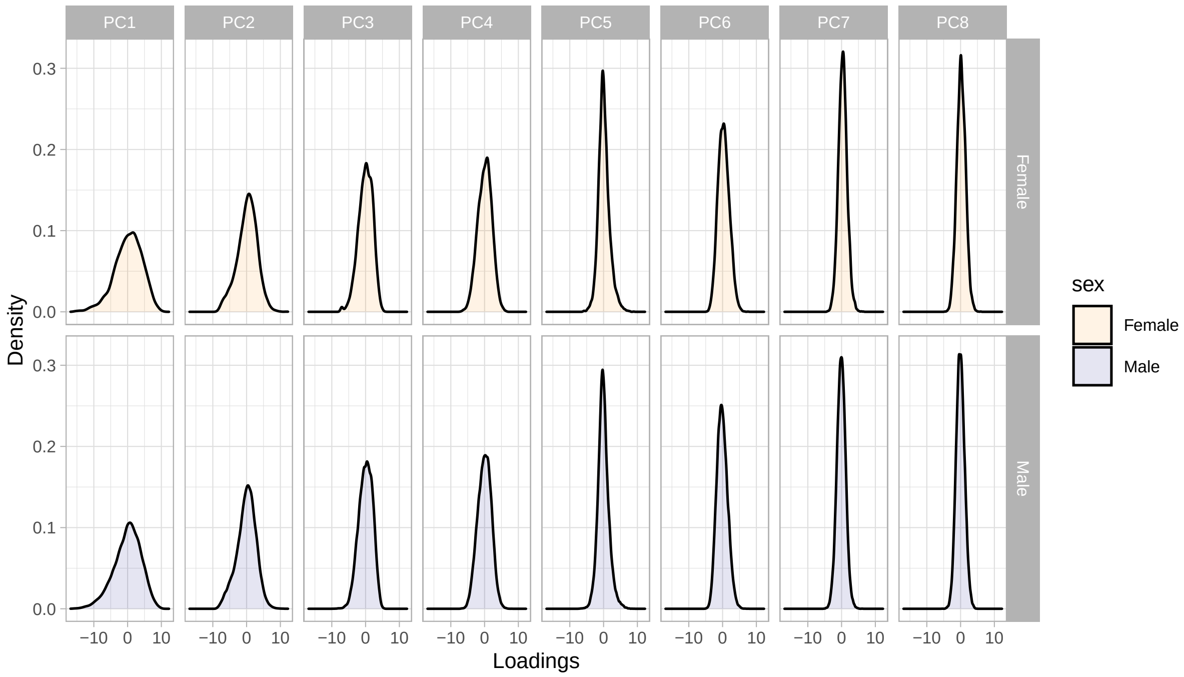

**Supp. Figure 2: FA loadings by sex.**

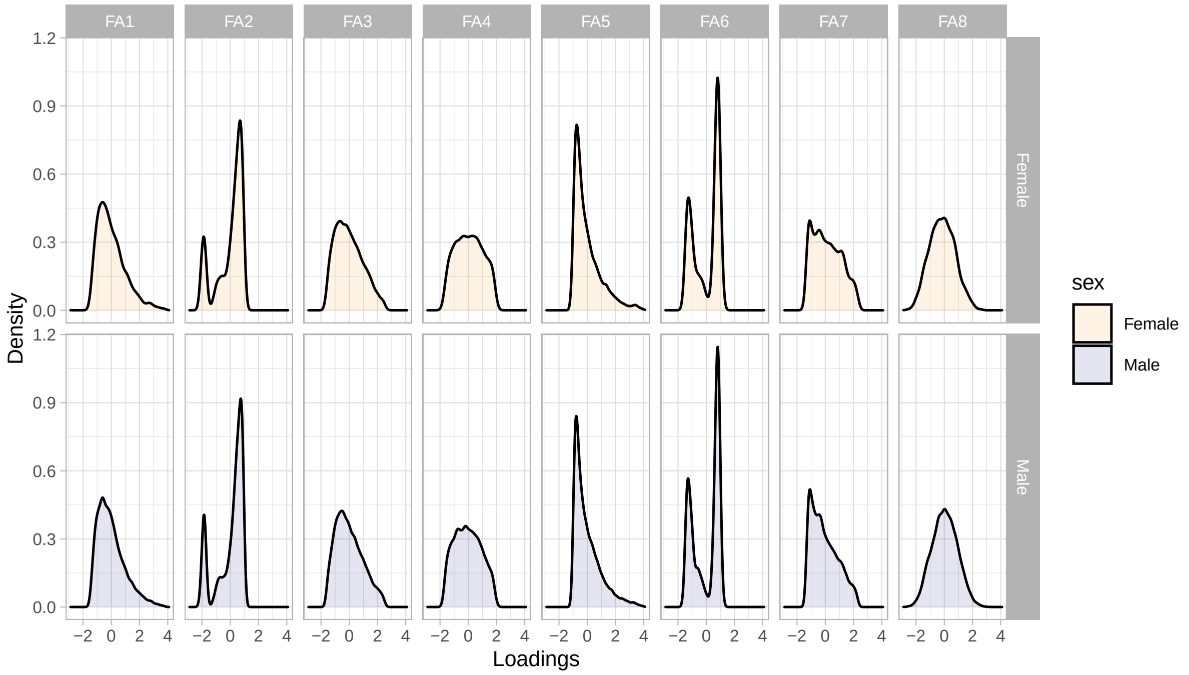


**Supp. Figure 3: VAE latent dimensions by sex.**

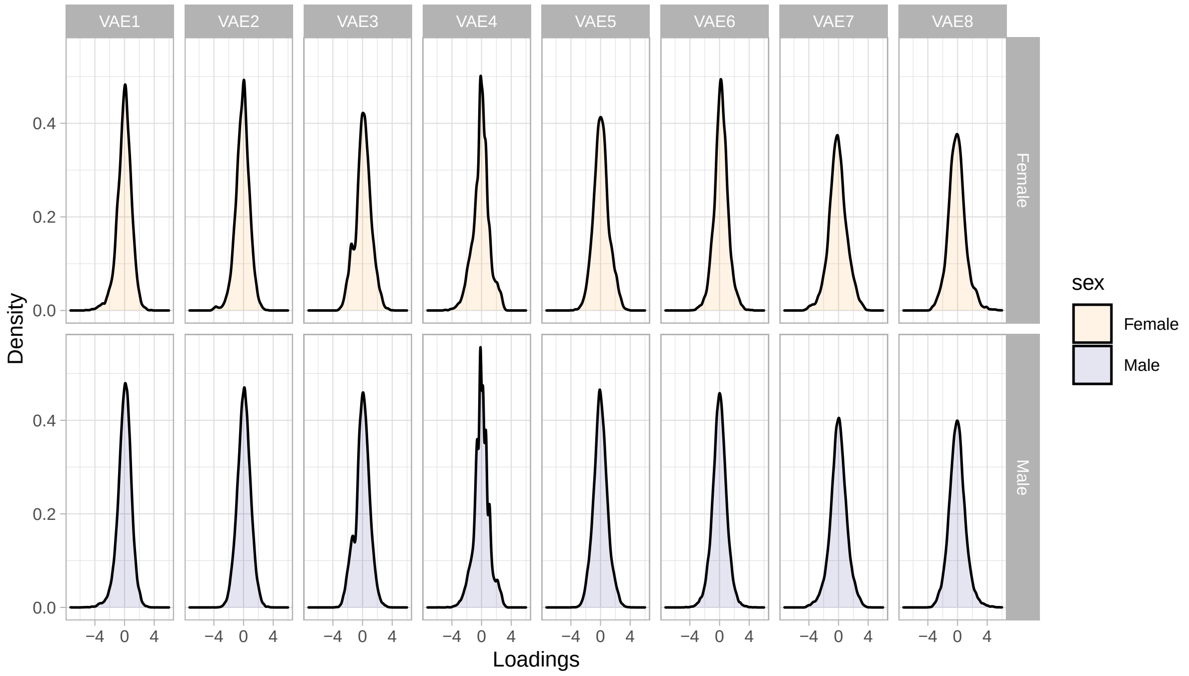

**Supp. Figure 4: Transformer latent dimensions by sex.**

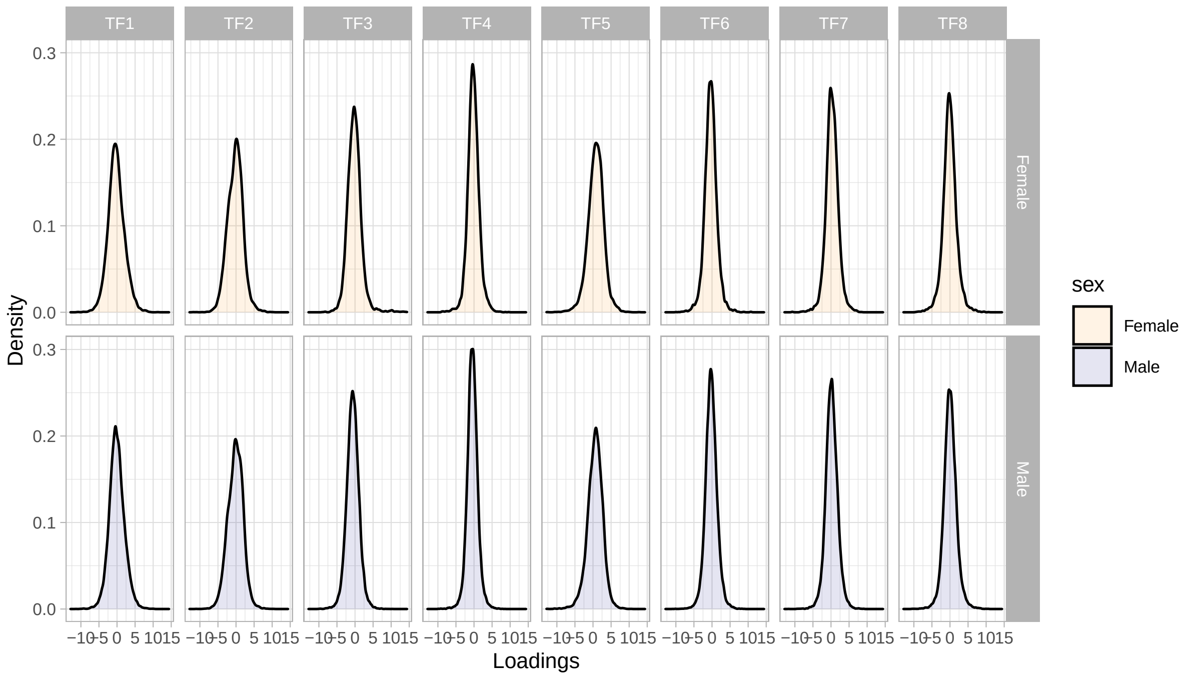


**Supp. Figure 5: Heritability of psychometric scales.**

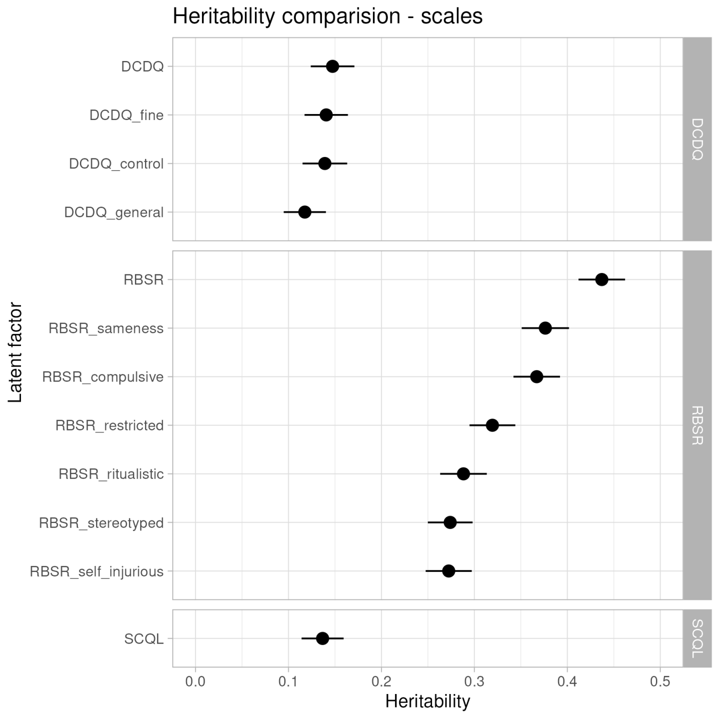

**Supp. Figure 6: Latent factor genetic correlation.**

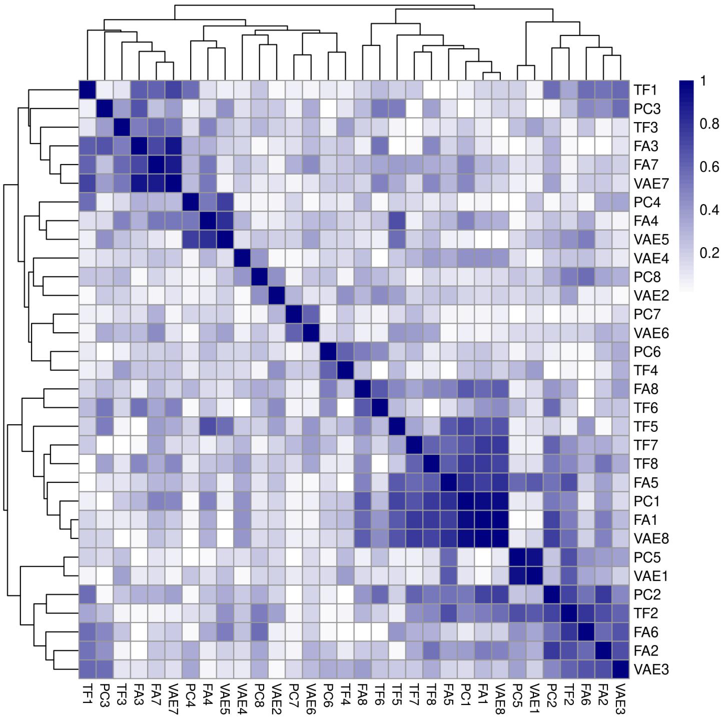


**Supp. Figure 7: Factor analysis loadings.**

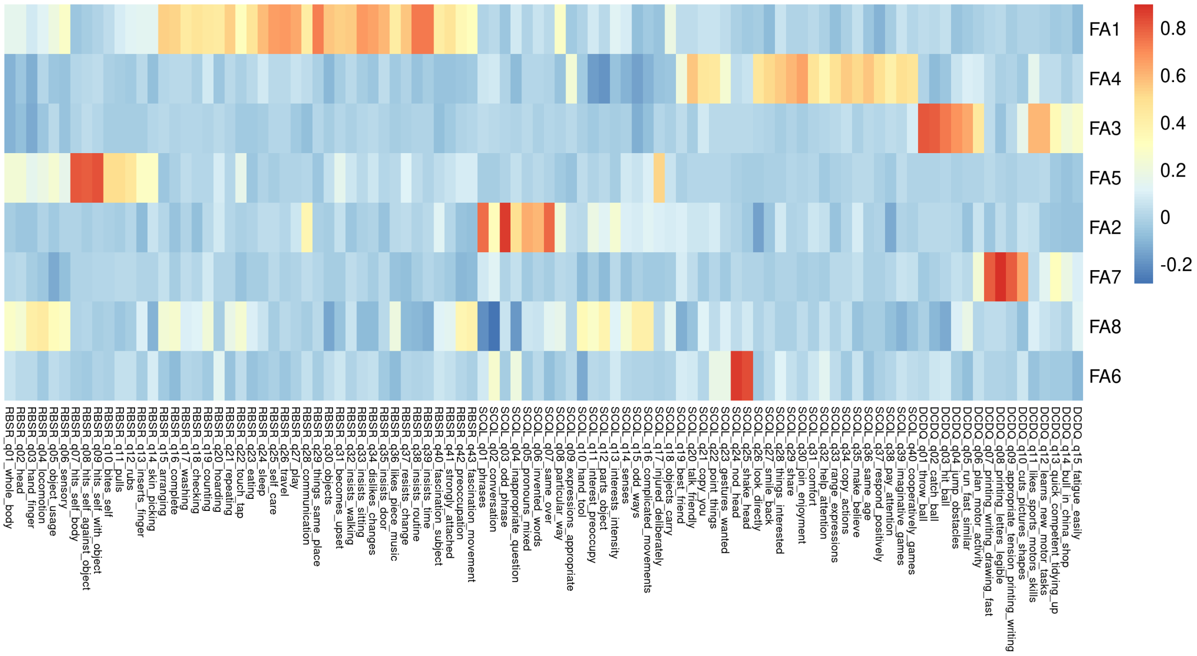

**Supp. Figure 8: VAE latent vector clinical correlations.**

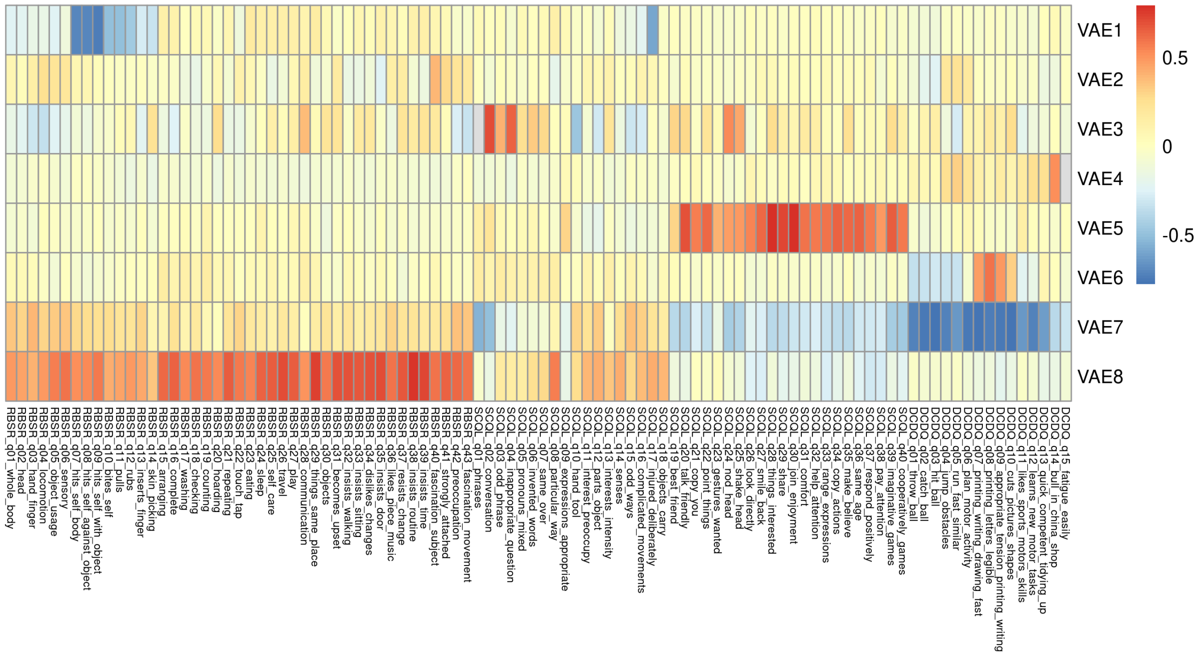

### Supplementary tables

#### Supp. Table 1: Genotypes excluded from study

| Genetic test | Subject count |
|---|---|
| 15q duplication | 166 |
| Trisomy 21 (Down Syndrome) | 119 |
| 22q11 deletion | 63 |
| 22q13_deletion | 63 |
| 15q deletion | 61 |
| 7q11.23 deletion | 44 |
| 16p11.2 deletion | 34 |
| 16p11.2 duplication | 28 |
| 1q21.1 duplication | 28 |
| 2p16.3 deletion | 19 |
| 22q11 duplication | 17 |
| XXY (Klinefelter syndrome) | 17 |
| 17q12 duplication | 15 |
| 3q29 duplication | 12 |
| 17q12 deletion | 11 |
| 1q21.1 deletion | 11 |
| Prader-Willi Syndrome | 11 |
| 16p12.2 deletion | 10 |
| 16p13.11 deletion | 10 |
| 17p13.3 duplication | 9 |
| 16p11.2 (unspecified) | 7 |
| 15q (unspecified) | 6 |
| 16p13.3 deletion | 5 |
| 3q29 deletion | 5 |
| 7q11.23 (unspecified) | 4 |
| 7q11.23 duplication | 4 |
| Xq28 duplication | 4 |
| 17q12 (unspecified) | 3 |
| 1q21.1 (unspecified) | 2 |
| 22q11 (unspecified) | 2 |
| 2q37.3 deletion | 2 |
| 17p11.2 duplication | 1 |
| 2p16.3 (unspecified) | 1 |
| 3q29 (unspecified) | 1 |
| 5q35 duplication | 1 |
| 6q16 deletion | 1 |
| XXYY | 1 |

**Supp. Table 2: SNPs with genome-wide significance.**

| Model | Factor | Chr | Pos | Lead SNP | Allele | P | Beta | SE |
|---|---|---|---|---|---|---|---|---|
| FA | FA1 | 1 | 3552525 | 1_3552525_G_A | A | 4.4e-08 | 0.145 | 0.026 |
| FA | FA1 | 1 | 3553817 | 1_3553817_T_C | C | 1.5e-08 | 0.152 | 0.027 |
| FA | FA1 | 2 | 210974349 | 2_210974349_G_T | T | 2.4e-08 | 0.223 | 0.040 |
| FA | FA1 | 3 | 114738193 | 3_114738193_T_C | C | 1.2e-08 | 0.328 | 0.057 |
| FA | FA1 | 4 | 46167100 | 4_46167100_GC_G | G | 1.0e-09 | -0.125 | 0.020 |
| FA | FA1 | 4 | 99322153 | 4_99322153_G_A | G | 1.5e-08 | 0.071 | 0.013 |
| FA | FA1 | 4 | 99322288 | 4_99322288_T_G | T | 4.0e-09 | 0.074 | 0.013 |
| FA | FA1 | 4 | 129413093 | 4_129413093_TTG_T | T | 4.7e-08 | 0.071 | 0.013 |
| FA | FA1 | 4 | 177571470 | 4_177571470_A_G | G | 1.6e-08 | 0.247 | 0.044 |
| FA | FA1 | 5 | 57755770 | 5_57755770_C_G | G | 2.1e-08 | -0.070 | 0.013 |
| FA | FA1 | 6 | 62858830 | 6_62858830_CA_C | CA | 4.3e-08 | 0.069 | 0.013 |
| FA | FA1 | 8 | 41855161 | 8_41855161_G_A | A | 3.0e-08 | 0.227 | 0.041 |
| FA | FA1 | 8 | 107058037 | 8_107058037_C_T | T | 4.9e-08 | -0.351 | 0.064 |
| FA | FA1 | 8 | 107065417 | 8_107065417_T_C | C | 3.0e-08 | -0.357 | 0.064 |
| FA | FA1 | 9 | 15148270 | 9_15148270_C_CT | CT | 2.0e-08 | 0.147 | 0.026 |
| FA | FA1 | 11 | 93326188 | 11_93326188_T_C | C | 1.3e-08 | 0.285 | 0.050 |
| FA | FA1 | 11 | 93344577 | 11_93344577_A_G | G | 4.3e-08 | 0.274 | 0.050 |
| FA | FA1 | 15 | 53683207 | 15_53683207_G_T | T | 1.5e-08 | 0.076 | 0.013 |
| FA | FA1 | 15 | 53684307 | 15_53684307_C_T | T | 1.4e-08 | 0.076 | 0.013 |
| FA | FA1 | 15 | 53684653 | 15_53684653_A_C | C | 7.0e-09 | 0.078 | 0.013 |
| FA | FA1 | 15 | 53685729 | 15_53685729_C_G | G | 3.0e-09 | 0.079 | 0.013 |
| FA | FA1 | 15 | 53691620 | 15_53691620_C(AGAT)×6_C | C | 1.3e-08 | 0.089 | 0.016 |
| FA | FA1 | 15 | 53691648 | 15_53691648_T_C | C | 9.0e-09 | 0.079 | 0.014 |
| FA | FA1 | 15 | 53691652 | 15_53691652_T_C | C | 3.0e-09 | 0.079 | 0.013 |
| FA | FA1 | 15 | 53691656 | 15_53691656_T_C | C | 1.3e-08 | 0.076 | 0.013 |
| FA | FA1 | 15 | 53691660 | 15_53691660_T_C | C | 1.9e-08 | 0.075 | 0.013 |
| FA | FA1 | 15 | 53691664 | 15_53691664_T_C | C | 1.9e-08 | 0.075 | 0.013 |
| FA | FA1 | 15 | 53691668 | 15_53691668_T_C | C | 2.5e-08 | 0.093 | 0.017 |
| FA | FA1 | 15 | 53693253 | 15_53693253_A_C | C | 1.4e-08 | 0.076 | 0.013 |
| FA | FA1 | 15 | 53695976 | 15_53695976_C_T | T | 7.0e-09 | 0.078 | 0.013 |
| FA | FA1 | 15 | 53703323 | 15_53703323_C_A | A | 5.0e-09 | 0.078 | 0.013 |
| FA | FA1 | 15 | 53704596 | 15_53704596_A_G | G | 1.3e-08 | 0.075 | 0.013 |
| FA | FA1 | 20 | 42735189 | 20_42735189_C_T | C | 4.8e-08 | 0.075 | 0.014 |
| FA | FA1 | 20 | 42735872 | 20_42735872_C_T | C | 1.3e-08 | 0.077 | 0.014 |
| FA | FA1 | 22 | 41202352 | 22_41202352_T_C | T | 1.2e-08 | 0.560 | 0.098 |
| FA | FA2 | 2 | 109474702 | 2_109474702_G_A | A | 3.3e-08 | -0.241 | 0.044 |
| FA | FA2 | X | 129859450 | 23_129859450_T_C | C | 4.4e-08 | -0.203 | 0.037 |
| FA | FA2 | X | 129859458 | 23_129859458_C_T | T | 4.4e-08 | -0.203 | 0.037 |
| FA | FA7 | 1 | 60797625 | 1_60797625_C_T | T | 3.6e-08 | 0.122 | 0.022 |
| FA | FA7 | 17 | 3268608 | 17_3268608_G_A | A | 2.8e-08 | 0.172 | 0.031 |
| FA | FA7 | 17 | 3275656 | 17_3275656_G_A | A | 1.7e-08 | 0.174 | 0.031 |
| FA | FA7 | 17 | 3399351 | 17_3399351_G_GAAAC | GAAAC | 2.0e-08 | 0.178 | 0.032 |
| VAE | VAE7 | 17 | 3268608 | 17_3268608_G_A | A | 2.4e-08 | -0.212 | 0.038 |
| VAE | VAE7 | 17 | 3275656 | 17_3275656_G_A | A | 2.3e-08 | -0.212 | 0.038 |
| VAE | VAE7 | 17 | 3371551 | 17_3371551_G_A | A | 4.9e-08 | -0.209 | 0.038 |
| VAE | VAE7 | 17 | 3397984 | 17_3397984_C_T | T | 4.5e-08 | -0.209 | 0.038 |
| VAE | VAE7 | 17 | 3399351 | 17_3399351_G_GAAAC | GAAAC | 2.9e-08 | -0.216 | 0.039 |

(Long SNP ids were shortened for increased readability.)

**Supp. Table 3: FA1 enriched Hallmark categories**

| Mechanism | setSize | enrichmentScore | NES | pvalue |
|---|---|---|---|---|
| KRAS SIGNALING DN | 151 | 0.718 | 1.190 | 0.012 |
| IL2 STAT5 SIGNALING | 164 | 0.719 | 1.194 | 0.012 |
| MITOTIC SPINDLE | 175 | 0.695 | 1.154 | 0.036 |
| E2F TARGETS | 153 | 0.696 | 1.153 | 0.040 |

**Supp. Table 4: FA2 enriched Hallmark categories**

| Mechanism | setSize | enrichmentScore | NES | pvalue |
|---|---|---|---|---|
| HYPOXIA | 159 | 0.688 | 1.218 | 0.022 |
| SPERMATOGENESIS | 104 | 0.698 | 1.236 | 0.027 |
| HEDGEHOG SIGNALING | 32 | 0.772 | 1.364 | 0.046 |
| ESTROGEN RESPONSE EARLY | 174 | 0.659 | 1.168 | 0.046 |

**Supp. Table 5: FA3 enriched Hallmark categories**

| Mechanism | setSize | enrichmentScore | NES | pvalue |
|---|---|---|---|---|
| MYC TARGETS V2 | 44 | 0.71 | 1.383 | 0.04 |

**Supp. Table 6: FA4 enriched Hallmark categories**

| Mechanism | setSize | enrichmentScore | NES | pvalue |
|---|---|---|---|---|
| XENOBIOTIC METABOLISM | 176 | 0.651 | 1.262 | 0.012 |
| NOTCH SIGNALING | 22 | 0.770 | 1.466 | 0.037 |

**Supp. Table 7: FA6 enriched Hallmark categories**

| Mechanism | setSize | enrichmentScore | NES | pvalue |
|---|---|---|---|---|
| TGF BETA SIGNALING | 41 | 0.773 | 1.462 | 0.007 |
| GLYCOLYSIS | 176 | 0.639 | 1.240 | 0.015 |
| BILE ACID METABOLISM | 101 | 0.650 | 1.248 | 0.045 |

**Supp. Table 8: FA7 enriched Hallmark categories**

| Mechanism | setSize | enrichmentScore | NES | pvalue |
|---|---|---|---|---|
| PI3K AKT MTOR SIGNALING | 87 | 0.627 | 1.274 | 0.048 |

**Supp. Table 9: FA8 enriched Hallmark categories**

| Mechanism | setSize | enrichmentScore | NES | pvalue |
|---|---|---|---|---|
| WNT BETA CATENIN SIGNALING | 35 | 0.748 | 1.471 | 0.025 |
| E2F TARGETS | 153 | 0.621 | 1.245 | 0.032 |

**Supp. Table 10: VAE1 enriched Hallmark categories**

| Mechanism | setSize | enrichmentScore | NES | pvalue |
|---|---|---|---|---|
| OXIDATIVE PHOSPHORYLATION | 173 | 0.613 | 1.262 | 0.021 |
| TGF BETA SIGNALING | 41 | 0.692 | 1.396 | 0.038 |
| ESTROGEN RESPONSE LATE | 174 | 0.597 | 1.228 | 0.039 |

**Supp. Table 11: VAE2 enriched Hallmark categories**

| Mechanism | setSize | enrichmentScore | NES | pvalue |
|---|---|---|---|---|
| APICAL JUNCTION | 172 | 0.648 | 1.307 | 0.004 |
| INTERFERON GAMMA RESPONSE | 169 | 0.602 | 1.215 | 0.032 |
| HEME METABOLISM | 170 | 0.593 | 1.195 | 0.042 |

**Supp. Table 12: VAE3 enriched Hallmark categories**

| Mechanism | setSize | enrichmentScore | NES | pvalue |
|---|---|---|---|---|
| IL2 STAT5 SIGNALING | 164 | 0.661 | 1.294 | 0.010 |
| REACTIVE OXYGEN SPECIES PATHWAY | 43 | 0.727 | 1.404 | 0.036 |
| APOPTOSIS | 133 | 0.653 | 1.274 | 0.037 |
| BILE ACID METABOLISM | 101 | 0.654 | 1.275 | 0.048 |

**Supp. Table 13: VAE4 enriched Hallmark categories**

| Mechanism | setSize | enrichmentScore | NES | pvalue |
|---|---|---|---|---|
| EPITHELIAL MESENCHYMAL TRANSITION | 173 | 0.645 | 1.267 | 0.016 |
| REACTIVE OXYGEN SPECIES PATHWAY | 43 | 0.742 | 1.426 | 0.017 |
| KRAS SIGNALING UP | 162 | 0.629 | 1.233 | 0.025 |

**Supp. Table 14: VAE5 enriched Hallmark categories**

| Mechanism | setSize | enrichmentScore | NES | pvalue |
|---|---|---|---|---|
| MYC TARGETS V2 | 44 | 0.752 | 1.408 | 0.013 |
| APICAL SURFACE | 41 | 0.746 | 1.391 | 0.014 |
| HEDGEHOG SIGNALING | 32 | 0.762 | 1.420 | 0.019 |
| XENOBIOTIC METABOLISM | 176 | 0.633 | 1.194 | 0.029 |
| INTERFERON GAMMA RESPONSE | 169 | 0.629 | 1.186 | 0.039 |
| COMPLEMENT | 167 | 0.625 | 1.178 | 0.043 |

**Supp. Table 15: VAE6 enriched Hallmark categories**

| Mechanism | setSize | enrichmentScore | NES | pvalue |
|---|---|---|---|---|
| GLYCOLYSIS | 176 | 0.634 | 1.274 | 0.009 |
| APICAL JUNCTION | 172 | 0.625 | 1.255 | 0.021 |

**Supp. Table 16: VAE8 enriched Hallmark categories**

| Mechanism | setSize | enrichmentScore | NES | pvalue |
|---|---|---|---|---|
| IL2 STAT5 SIGNALING | 164 | 0.681 | 1.334 | 0.002 |
| KRAS SIGNALING DN | 151 | 0.643 | 1.258 | 0.014 |
